# Overlooked *Candida glabrata* petites are echinocandin tolerant, induce host inflammatory responses, and display poor *in vivo* fitness

**DOI:** 10.1101/2023.06.15.545195

**Authors:** Amir Arastehfar, Farnaz Daneshnia, Hrant Hovhannisyan, Diego Fuentes, Nathaly Cabrera, Christopher Quintin, Macit Ilkit, Nevzat Ünal, Suleyha Hilmioğlu-Polat, Kauser Jabeen, Sadaf Zaka, Jigar V. Desai, Cornelia Lass-Flörl, Erika Shor, Toni Gabaldon, David S. Perlin

## Abstract

Small colony variants (SCVs) are relatively common among some bacterial species and are associated with poor prognosis and recalcitrant infections. Similarly, *Candida glabrata* – a major intracellular fungal pathogen – produces small and slow-growing respiratory-deficient colonies, termed “petite.” Despite reports of clinical petite *C*. *glabrata* strains, our understanding of petite behavior in the host remains obscure. Moreover, controversies exist regarding in-host petite fitness and its clinical relevance. Herein, we employed whole-genome sequencing (WGS), dual-RNAseq, and extensive *ex vivo* and *in vivo* studies to fill this knowledge gap. WGS identified multiple petite-specific mutations in nuclear and mitochondrially-encoded genes. Consistent with dual-RNAseq data, petite *C*. *glabrata* cells did not replicate inside host macrophages and were outcompeted by their non-petite parents in macrophages and in gut colonization and systemic infection mouse models. The intracellular petites showed hallmarks of drug tolerance and were relatively insensitive to the fungicidal activity of echinocandin drugs. Petite-infected macrophages exhibited a pro-inflammatory and type I IFN-skewed transcriptional program. Interrogation of international *C*. *glabrata* blood isolates (*n*=1000) showed that petite prevalence varies by country, albeit at an overall low prevalence (0–3.5%). Collectively, our study sheds new light on the genetic basis, drug susceptibility, clinical prevalence, and host-pathogen responses of a clinically overlooked phenotype in a major fungal pathogen.

**Importance:** *Candida glabrata* is a major fungal pathogen, which is able to lose mitochondria and form small and slow-growing colonies, called “petite”. This attenuated growth rate has created controversies and questioned the clinical importance of petiteness. Herein, we have employed multiple omics technologies and in vivo mouse models to critically assess the clinical importance of petite phenotype. Our WGS identifies multiple genes potentially underpinning petite phenotype. Interestingly, petite *C. glabrata* cells engulfed by macrophages are dormant and therefore are not killed by the frontline antifungal drugs. Interestingly, macrophages infected with petite cells mount distinct transcriptomic responses. Consistent with our ex-vivo observations, mitochondrial-proficient parental strains outcompete petites during systemic and gut colonization. Retrospective examination of *C. glabrata* isolates identified petite prevalence a rare entity, can significantly vary from country to country. Collectively, our study overcomes the existing controversies and provides novel insights regarding the clinical relevance of petite *C. glabrata* isolates.

## Introduction

One of the strategies used by microbes to rapidly adapt and survive in stressful conditions is to fine-tune central carbon metabolism to achieve phenotypic plasticity (1–7). A remarkable example of this is the occurrence of small colony variant (SCV) bacterial isolates following antibiotic exposure, most notably in *Staphylococcus aureus*, which are implicated in antibiotic therapeutic failure, difficult-to-treat recurrent infections, and high disease severity (1, 4, 8). Strikingly, SCVs are effectively phagocytosed by host cells, and transcriptomic studies have shown that SCVs do not elicit potent immune responses or damage host cells (5, 9). Accordingly, it is believed that low virulence and slow growth are strategies allowing SCVs to successfully exploit the host cells they infect, avoiding the cytotoxic action of the immune system and protecting themselves from direct exposure to lethal antibiotics. Upon cessation of antibiotic treatment, however, such SCVs can rapidly revert to the fully virulent wild-type (WT) phenotype, causing relapse and seeding chronic infections (5, 9).

Despite being an essential organelle in most eukaryotes, some eukaryotic species, such as baker’s yeast *Saccharomyces cerevisiae*, can lose mitochondria or oxidative respiratory functions under certain conditions, and the resultant cells, known as petite mutants, are viable, forming small, slow-growing colonies that resemble SCVs (10, 11). The appearance of petite mutants has also been noted in clinical samples obtained from patients infected with *Candida glabrata* (2, 12), a prominent human fungal pathogen more closely related to *S*. *cerevisiae* than to *C*. *albicans* (13). Although petite clinical *C*. *glabrata* isolates were previously thought to be rare, a recent study discovered that approximately 11% of examined clinical *C*. *glabrata* isolates (16/146) displayed hallmarks of petiteness, including small and slow-growing colonies, lack of mitochondrial membrane potential, and inability to grow on non-fermentable carbon sources (7). Interestingly, petite *C*. *glabrata* shows resistance to fungistatic azole drugs due to overexpression of ABC transporters (*CDR1*, *CDR2*, and *SNQ2*) and their transcriptional regulator (*PDR1*), in contrast to the canonical azole resistance mechanism driven by gain-of-function *PDR1* mutations (2, 3, 7, 12, 14, 15). However, how petites are affected by fungicidal drugs (echinocandins and polyenes) has not been determined.

Although *C*. *glabrata* can be engulfed by macrophages, it is highly resistant to macrophage-mediated killing and can survive and proliferate within these host cells (16). Interestingly, similar to SCVs, petite cells are more effectively phagocytosed relative to non-petite strains (7). However, it is not known whether petite strains affect the host phagocytic cells they infect differently than non-petite strains, and vice versa. Furthermore, their effect in the host has remained controversial, with some studies reporting substantially higher mortality and fungal burdens in petite strain-infected mice (14), whereas other studies have found otherwise (15, 17). These differences may be partly due to the origin of petite isolates or their underlying genetic basis. In *S*. *cerevisiae*, petiteness can be caused by different genetic mutations affecting mitochondrial biogenesis or function, but whether this is true for *C*. *glabrata* petites is unknown. One study focused on the *C*. *glabrata* gene encoding a mitochondrial DNA polymerase, *MIP1*, and found the same *MIP1* SNPs in both petite and non-petite strains (7), indicating that other mutations can induce a petite phenotype.

Herein, we used clinical and laboratory-generated petite isolates to systematically address these questions. Whole genome sequencing revealed multiple genetic mechanisms underlying the petite phenotype, including mutations affecting proteins involved in mitochondrial mRNA stability. Our dual RNA-seq analysis of *C*. *glabrata*-infected macrophages indicated that intracellular petite cells showed signatures of non-growth and that petite-infected macrophages exhibited pronounced type-I interferon and proinflammatory cytokine transcriptional responses at later infection stages compared to macrophages infected with non-petite strains. We also discovered that petites of different biological origins (clinical or laboratory-derived) are readily outcompeted by their non-petite parental strains in macrophage interaction assays, as well as in gut colonization and systemic infection mouse models. However, the petite strains showed a fitness advantage over non-petite strains during echinocandin treatment. Finally, we assessed the prevalence of petites in a large (1,000 strains) international collection of *C*. *glabrata* blood isolates, and although petites were extremely rare (9/1,000, 0.9%), their prevalence varied depending on the geography (0–3.3% of the total blood isolates). Additionally, most of the petite isolates recovered (89%) were a mixture of large and small colonies reminiscent of SCVs. Altogether, our paper sheds new light on the genetic basis of petiteness in *C*. *glabrata*, its clinical prevalence, and its potential implications for infection and drug resistance.

## Results

### Petite isolate collection, characterization, and evaluation of metabolic deficiencies

We studied a collection of strains comprising clinical and laboratory-derived petite *C*. *glabrata* isolates (Supplementary Table 1). BYP40 is a previously reported non-petite isolate recovered from the blood of an azole-naïve patient five days after hospitalization, and BYP41 is a petite isolate recovered from the same patient after fluconazole treatment (12). Interestingly, the patient infected with BYP41 did not respond to fluconazole, and the bloodstream infection caused by this isolate was ultimately cleared by amphotericin B (12). We also identified a petite *C*. *glabrata* isolate in our laboratory collection (DPL248), for which we did not have clinical data or the parental strain. Finally, we obtained four laboratory-derived petite strains, namely, C5, D5, F2, and G5, by evolving the *C*. *glabrata* type strain CBS138 in the presence of fluconazole (see Figures S1A and S1B and Methods section). Briefly, CBS138 was inoculated in RPMI containing 64 µg/ml of fluconazole, and fluconazole-resistant (FLZR) colonies (MIC ≥64 µg/ml) lacking *PDR1* mutations were selected and analyzed further for petite traits. The final four independently derived petite isolates were unable to grow on YP-glycerol (YPG) agar plates, were FLZR (Figure 1A), overexpressed *PDR1*, *CDR1*, *CDR2*, and *SNQ2* (Figure 1B), and had significantly lower ATP levels (Figure 1C) and mitochondrial membrane potential (Figure 1D) compared to non-petite isolates.

**Figure 1.**
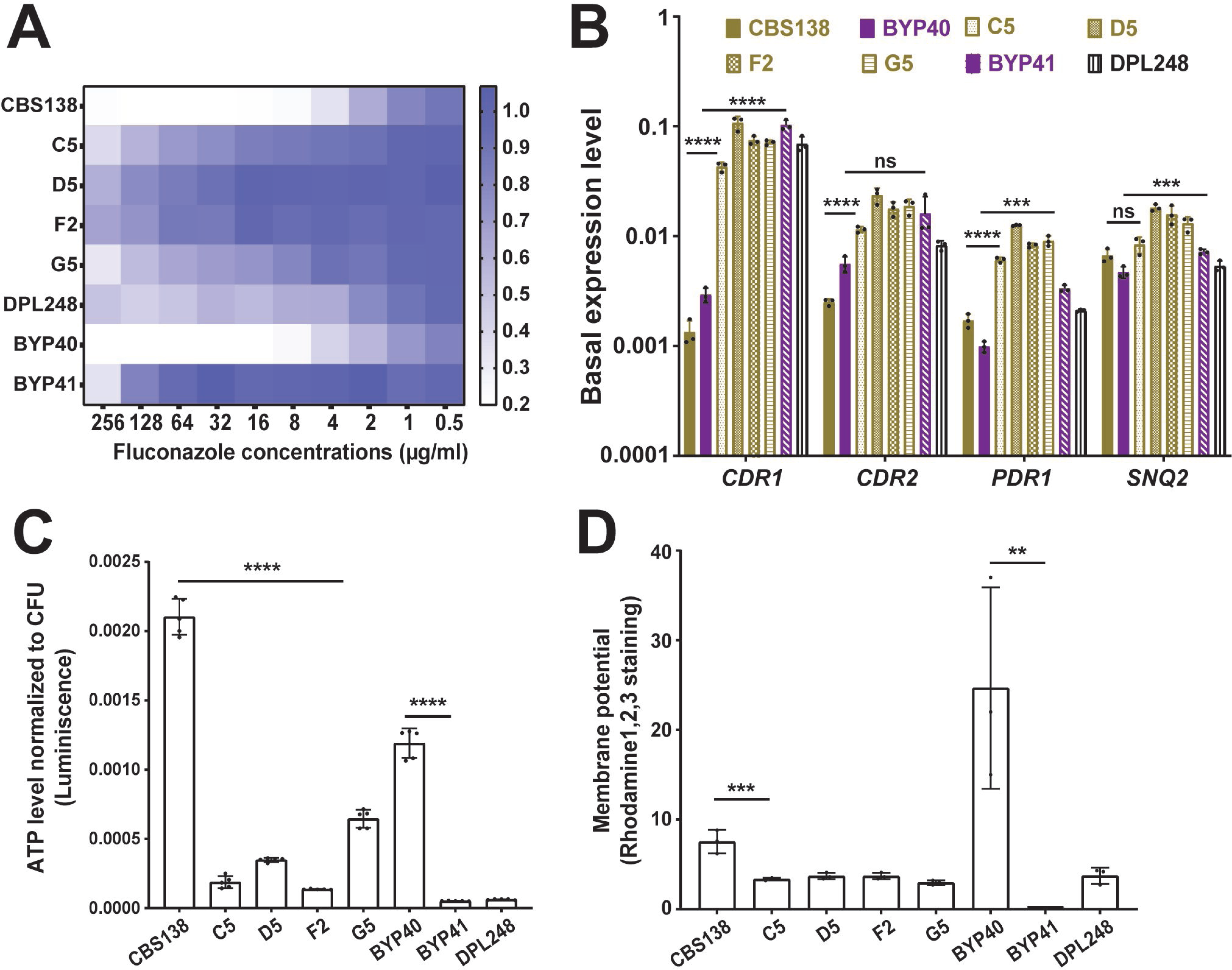
Characteristics of petite *C. glabrata* isolates. All petite isolates included in this study were fluconazole resistant (A) and did not carry *PDR1* gain-of-function mutations, as confirmed by *PDR1* sequencing. Petite isolates have a higher basal level of *PDR1* and efflux pumps under its control (*CDR1*, *CDR2*, and *SNQ2*) once measured by real-time PCR (3 biological replicates, ***<0.01, ****<0.001, two-tailed t-test) (B). Petites had a lower ATP level (5 biological replicates, ****<0.001, two-tailed t-test) (C) and mitochondrial membrane potential (3 biological replicates, ***<0.01 and **=0.01, two-tailed t-test) (D) than their respiratory proficient counterparts. ATP: Adenosine triphosphate.

Because mitochondria are involved in and influence multiple biosynthetic processes, including amino acid, heme, and nucleotide production (18, 19), petite mutants may be deficient in certain metabolites. Indeed, petite mutants of *S*. *cerevisiae* exhibit deficiencies in leucine, arginine, glutamate, and glutamine but not in amino acids derived from the reductive part of the TCA cycle, such as aspartate (10). In general, with the exception of G5, the growth of petite isolates was significantly improved upon supplementation with arginine, leucine, adenine, and thymidine (Figures S1C and S1D), which is similar to *S*. *cerevisiae* petites (10). However, unlike in *S*. *cerevisiae*, *C*. *glabrata* petites’ slow growth was not improved with glutamate supplementation, whereas glutamine and hemin improved the growth rate of both petites and non-petite isolates similarly. These observations confirm that the majority of petite *C*. *glabrata* strains exhibit the metabolic deficiencies expected of cells with non-functional mitochondria.

Finally, we assessed the stability of the petite phenotype by passaging the petite isolates up to 30 times in YPD (overnight cultures) and streaking multiple colonies on YPG plates to look for non-petite revertants. Whereas the G5 strain readily reverted to a non-petite phenotype after the second passage, the rest of the CBS138-derived and clinical petite isolates were stable, and no revertants were observed after 30 passages. Thus, most of the petites were stable, whereas G5, which also showed the fastest growth rate as well as higher ATP levels among the petite strains, was reversible.

### Whole genome sequencing reveals mutations associated with petite strains

Loss-of-function mutations in mitochondrial DNA polymerase Mip1 have been shown to result in a petite phenotype in *C*. *glabrata* (7). We sequenced the *MIP1* gene of petites and their non-petite parents (Supplementary Table 1) and found that the petites did not harbor any mutations absent in their parental strains and that *MIP1* polymorphisms predicted the sequence type of sequenced isolates (see Figure 2A and Supplementary Table 1 indicating that mutations elsewhere in the genome caused the petite phenotype. To identify these mutations, we used Illumina next-generation sequencing. Whereas DPL248 and petites derived from CBS138 had a mitochondrial genome coverage of 2x–6x of the nuclear chromosome, BYP41 had minimal mitochondrial DNA relative content (Figure S2A). We identified high confidence variants that were different between BYP40 and BYP41 (Supplementary Table 2) and those that were present in some, but not all, of the four CBS138-derived petite strains (Supplementary Table 2), as shared variants in these four strains were likely present in their non-petite parent. The lack of a close non-petite parent strain for DPL248 prevented us from identifying recently acquired mutations. To exclude polymorphisms due to phylogenetic distance across clades, we determined the MLST type from the DPL248 genomic sequence as ST7 and identified two previously sequenced strains, CST35 and EB0911Sto, as its closest sequenced relatives (20). We thus excluded variants between DPL248 and CBS138 that were also present in either of these two strains, resulting in a restricted list of DPL248 variants (Supplementary Table 2). Gene ontology enrichment analyses for genes present in the three tables identified pathways related to mitochondrial functions and components of the cell wall (Supplementary Table 3). Cell wall-encoding genes are known to be highly variable and are probably unrelated to the petite phenotype (20, 21).

**Figure 2.**
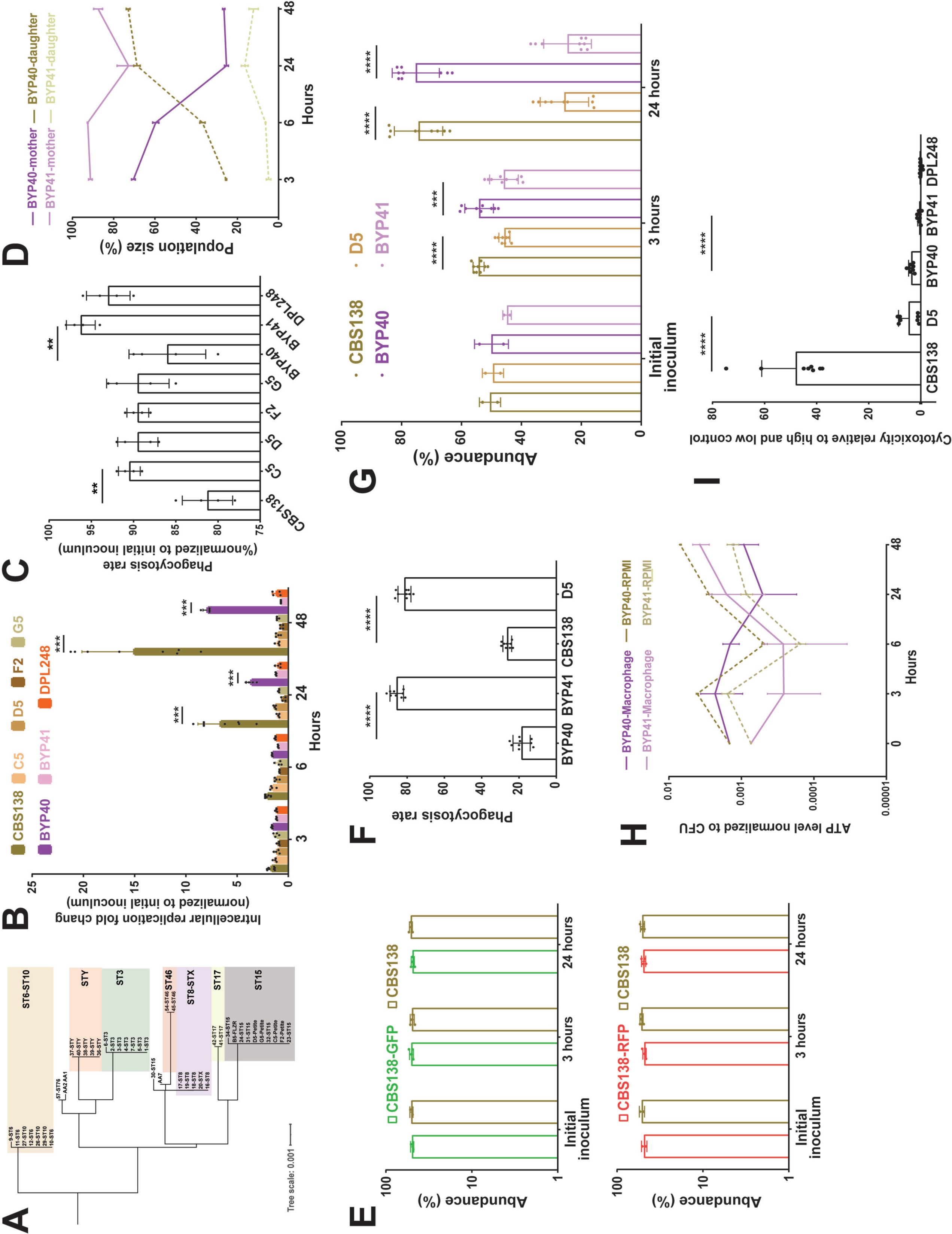
*MIP1* sequencing of diverse clinical isolates revealed that polymorphisms occurring in isolates better reflect the sequence type rather than petite phenotype (A). The interaction of petite and their respective parental isolates with THP1 macrophages. Macrophages infected with respective isolates and intracellular replication was measured 3, 6, 24, and 48 hours after infection and the data were normalized against the initial inoculum used to treat the macrophages. Petite isolates did not show intracellular replication, unlike their respiratory proficient isolates (4-8 biological replicates, ***<0.001, two-tailed t-test) (B), whereas petite isolates had a significantly higher rate of phagocytosis rate (4 biological replicates, **<0.01, two-tailed t-test) (C). Macrophages were infected with FITC-stained BYP40 and NYP41 and counterstained with AF647 after macrophage lysis at each timepoints and the single and double positive events were measured by fluorescent activated cell sorting (FACS). FITC staining and AF647 counterstaining of intracellular BYP40 and BYP41 revealed that petites have an extremely limited growth as all cells were double-stained, while BYP40 showed a significant intracellular growth as evidenced by a high proportion of single-stained cells (3 biological replicates) (D). Genomic interaction of GFP and RFP into CBS138 did not impact the intracellular growth and both transformants showed equally replicated inside the macrophages (4 biological replicates) (E). RFP-expressing petite isolates were effectively phagocytosed by THP1 macrophages (4 biological replicates, ****<0.00001, two-tailed t-test) (F), whereas they were outcompeted by their parental strains (9 biological replicates, ***<0.01 and ****<0.00001, two-tailed t-test) (G). Petite mutants enter a near dormant state after internalization by macrophages. The ATP level of BYP40 and BYP41 was normalized against CFU incubated in RPMI or macrophages at different time-points. Intracellular BYP41 had a significantly lower ATP level at early hours, whereas its ATP level was significantly higher than BYP40 48-hours (6 biological replicates) (H). Unlike their parental isolates, petite isolates imposed the least cytotoxicity 24-hours post-infection as measured by lactate dehydrogenase (8 biological replicates, ****<0.00001, two-tailed t-test) (I). FACS: fluorescent activated cell sorting.

To identify mutations underlying the petite phenotype, we identified genes harboring non-synonymous variants (not necessarily in the same position) in two or more of the above-mentioned tables. Three genes were selected as the most likely candidates to explain the petite phenotype across the petite strains (Supplementary Table 3), *COX3*, *SSU*, and *Cgai1*, which are all implicated in critical mitochondrial functions. Additionally, we identified three other genes associated with mitochondrial function that appeared uniquely in CBS138-derived petites, including *RDM9* (CAGL0F07469g, regulating mRNA stability, translational initiation in the mitochondrion), *MSY1* (CAGL0H05775g, involved in group I intron splicing, mitochondrial tyrosyl-tRNA aminoacylation and mitochondrion localization), and *CIT1* (CAGL0H03993g, encoding citrate of the mitochondrial tricarboxylic acid cycle). Additionally, two other mutations listed in Supplementary Table 3 that appear unique to DPL248 were related to mitochondrial functions, including mutations in *CaglfMp07* (*Cgai3*), a putative endonuclease encoded by the first three exons and part of the third intron (a group I intron) of the mitochondrial *COX1* gene, and *CaglfMt24* (M(CAU)9mt), a mitochondrial methionine tRNA with a CAU anticodon, one of two tRNA-Met encoded on the mitochondrial genome.

To understand the mechanisms underlying petite phenotype and given the complexity of mitochondrial genome manipulation, we focused only on nuclear encoded genes with known function, i.e., *CIT1*, *MSY1*, and *RDM9* and constructed the respective deletion mutants in a CBS138 background. Although *cit1Δ* and *rdm9Δ* deletions were readily generated, multiple attempts to delete *MSY1* failed. Interestingly, *rdm9Δ*, but not *cit1Δ*, could not grow on YPG agar plates and was FLZR (Figures S2B ). It significantly overexpressed *CDR1*, *CDR2*, *PDR1*, and *SNQ2* (Figure S2C) and had a significantly lower mitochondrial membrane potential and ATP level (Figures S3A and S3B). Moreover, similar to petite isolates, *rdm9Δ* poorly grew in YNB and showed leucine, arginine, glutamine, menadione, and thymidine dependency (Figures S3C and S3D). These results underscore the power of our comparative genomics approach used to discover genes potentially involved in petite phenotype and highlight the importance of mitochondrial mRNA stability in mitochondrial function. They also elucidated the complex nature of the genetic underpinning of petiteness, which precluded a straightforward association of a single genetic defect among strains.

### Intracellular petite cells are non-growing, dormant-like, and exert minimal damage to macrophages

Given the observed growth deficiencies in minimal media and that macrophages impose prominent carbon starvation (22–24), we reasoned that petite isolates should not grow inside macrophages. To test this hypothesis, we measured the replication of non-petite and petite *C*. *glabrata* isolates inside THP1 macrophages. After 3 hours, THP1 cells infected with *C*. *glabrata* isolates were extensively washed with PBS to remove non-adherent yeast cells and provided with fresh RPMI, and the intracellular replication rate was measured 3-, 6-, 24-, and 48- hours post-infection (pi) by plating and CFU counting. Moreover, non-adherent cells obtained at 3 hours were plated to measure the phagocytosis rate. Whereas non-petite isolates showed high levels of replication at 24- and 48- hours pi, petite counterparts did not exhibit intracellular growth (Figure 2B). In contrast, petites had a significantly higher phagocytosis rate than non-petite isolates, consistent with previous observations (7) (Figure 2C). Similar to petite isolates, *rdm9Δ* also did not show intracellular growth (Figure S3E).

To substantiate the observation that petites are unable to grow intracellularly, we infected THP1 macrophages with either non-petite or petite *C*. *glabrata* cells stained with fluorescein isothiocyanate (FITC), which is not transferred to the daughter cells. After releasing the intracellular *C*. *glabrata* cells at each time-point, they were counter-stained with Alexa Flour-647 Concanavalin A (AF-647), which stains both mother and daughter cells (7). Therefore, intracellular daughter yeast cells will be single positive for AF-647, whereas the mother cells will be double positive for FITC and AF-647, and their relative amounts could be measured via flow-cytometry (7). We chose BYP40 and BYP41 and monitored the abundance of mother and daughter yeast cells at 3-, 6-, 24-, and 48- hours pi (Figure 2D). Consistent with our previous experiments, the progressive intracellular replication of BYP40 was reflected by a significant increase in the proportion of daughter cells, whereas BYP41 showed a relatively stable proportion of daughter and mother cells throughout the course of the experiment, indicative of the absence of intracellular growth.

Next, we investigated whether petites could be outcompeted by their parental non-petite isolates inside the macrophages. Therefore, we constructed plasmid-borne DNA cassettes of green fluorescent protein (GFP) or red fluorescent protein (RFP) in close proximity to the nourseothricin N-acetyl transferase (NAT) gene and integrated these cassettes into chromosome F of petite and non-petite strains using CRISPR/Cas9 and nourseothricin selection. Notably, GFP and RFP mutants showed similar growth rates compared to CBS138 after 24 hours (Figure 2E). The phagocytosis rate was measured 3 hours pi, whereas the intracellular replication was assessed 3- and 24- hours pi. Consistent with our mono-culture experiments, petites had significantly higher phagocytosis rates (Figure 2F), and inside macrophages, they were outcompeted by their non-petite counterparts 24 hours pi (Figure 2G).

Since intracellular petites did not replicate inside macrophages, we wondered whether macrophage internalization triggers low metabolic activity. Measurement of ATP levels was used as a proxy for metabolic activity determination. The ATP levels of intracellular and planktonic BYP40 and BYP41 were determined at 3-, 6-, 24-, and 48 hours pi, and the values obtained were normalized against the corresponding colony forming units (CFU). As expected, the ATP levels of planktonic BYP41 were lower than those of BYP40 at all time-points (Figure 2H). Interestingly, the ATP level of intracellular BYP41 followed a dynamic trend, being extremely low at 3 and 6 hours and recovering and surpassing that of planktonic conditions at 48- hours. Given the lack of mitochondrial activity and the lack of fermentable carbon sources in the phagosome, such a surge in ATP level might be either an indication of acquiring ATP from the host (25) or likely glycolytic ATP production with reduced metabolic activity.

Given this observation and the lack of intracellular growth, we hypothesized that similar to SCVs, petites may be less cytotoxic toward THP1 macrophages than their non-petite counterparts. Cytotoxicity was investigated by assessing lactate dehydrogenase (LDH) levels 24 hours pi (26). Consistent with our hypothesis, the LDH levels of the THP1 macrophages infected with non-petite isolates were significantly higher than those of the petite counterparts (Figure 2I). Altogether, these findings suggest that unlike non-petite parental strains, petites do not grow inside THP1 macrophages, and this lack of growth is reflected by significantly lower cytotoxicity.

### Dual RNA-seq of petite and non-petite isolates

To further disentangle the differences between petite and non-petite *C*. *glabrata* and how the petite phenotype influences interactions with macrophages, we used a time-course dual RNA-seq approach (27, 28). THP1 macrophages were infected with either petite strains or their parental strains, and the infected macrophages were collected at 3 hr and 24 hr pi for RNA isolation and RNA-seq analysis.

First, we explored the transcriptional patterns of the fungal cells upon interaction with macrophages. The overall transcriptional profiles of infecting *C*. *glabrata* isolates (Figure 3A) show that the main differences between cell types are driven by the petite phenotype and, to a lesser extent, by the time-point of infection. Interestingly, when growing in RPMI, the studied strains showed larger differences between time points, although they still showed differences due to petiteness. These observations indicate that the stressful environment within macrophages exacerbates the transcriptomic divergence between petite and non-petite *C*. *glabrata*. We then aimed to shed light on functional differences due to petiteness among laboratory-derived (D5 *vs.* CBS138) and clinical isolates (BYP40 *vs.* BYP41). To do this, we performed differential gene expression analysis with subsequent aggregated Gene Ontology (GO) term enrichment analysis (Figure 3B). Interestingly and consistent with our *ex vivo* data, petite isolates downregulated processes associated with replication (Figure 3C).

**Figure 3.**
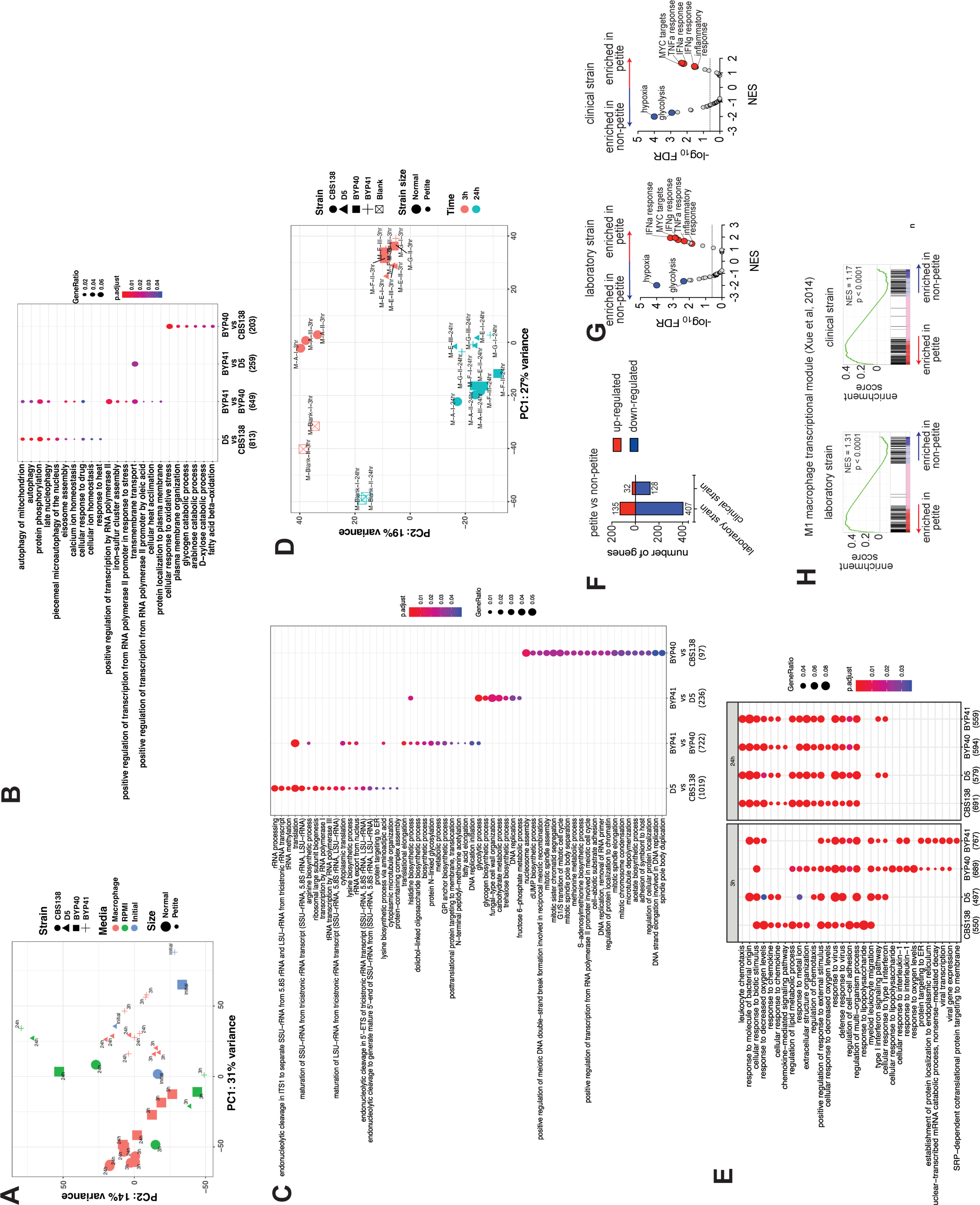
Principal Component Analysis (PCA) plot of all studied *C. glabrata* samples across studied conditions. The plot is based on vst-transformed read count data generated by DESeq2. Labels on the data points correspond to time points of the experiments. Percentages on PC1 and PC2 axes indicate the total amount of variance described by each axis (A). GO term enrichment analysis (category “Biological Process”) of up-regulated genes of C. glabrata at a given comparison shown on the X axis. The numbers underneath the comparisons correspond to the “counts” of clusterProfiler (i.e. total number of genes assigned to GO categories). GeneRatio corresponds to the ratio between the number of input genes assigned to a given GO category and “counts”. Only significant (padj<0.05) enrichments are shown. Adjustment of p-values is done by Benjamini-Hochberg procedure (B). Petitness-specific GO terms such as tRNA and rRNA related processes, biosynthesis of several amino acids as arginine and lysine, among others. When compared to laboratory-derived petites, clinical petites showed down-regulation of carbohydrate biosynthesis and fungal cell-wall related processes. Finally, the comparisons of non-clinical strains of normal size showed down-regulation of various processes in the clinical stains, such as nucleosome and mitotic spindle assembly, methionine and acetate metabolism, etc. (C). Principal Component Analysis (PCA) plot of all studied macrophage samples. The plot is based on vst-transformed read count data generated by DESeq2. Labels on the data points correspond to internal sample identifiers. Percentages on PC1 and PC2 axes indicate the total amount of variance described by each axis (D). GO term enrichment analysis (category “Biological Process”) of up-regulated genes of macrophages infected with *C. glabrata* strains (as depicted on X axis) compared to unchallenged macrophages (E). The numbers underneath the comparisons correspond to the “counts” of clusterProfiler (i.e. total number of genes assigned to GO categories). GeneRatio corresponds to the ratio between the number of input genes assigned to a given GO category and “counts”. Only significant (padj<0.05) enrichments are shown. Adjustment of p-values is done by Benjamini-Hochberg procedure. *C. glabrata* petite strains induce a pro-inflammatory transcriptional program in human THP-1 macrophage cells. Summary data of differentially expressed transcripts in THP-1 macrophages at 24h post-challenge; comparisons are between THP-1 transcriptomes challenged with petite vs. non-petite *C. glabrata* laboratory or clinical strains (F). gene set enrichment analysis (GSEA) indicating significantly enriched “Hallmark” pathways of the Molecular Signatures Database, based on the RNA-seq data from THP-1 cells at 24h post-fungal challenge. The pathways are displayed based on the normalized enrichment score (NES) and the false discovery rate (FDR). The dotted line marks an FDR value of 0.25, while a select top enriched pathways are indicated Blue (enriched in non-petite) and Red (enriched in Petite) (G). GSEA enrichment plots depicting enrichment of the M1 macrophage transcriptional module (Xue et al, 2014) comparing transcriptomes THP-1 cells challenged with the petite vs. non-petite C. glabrata laboratory or clinical strains, at 24h post-challenge (H). The p value reported here is the nominal p-value while NES is the normalized enrichment score.

Our GO analysis results show that many biological processes, such as autophagy of mitochondria, protein phosphorylation, calcium ion homeostasis, and eiosome assembly, are exclusively upregulated in petite mutants. On the other hand, we observed biological processes specifically up-regulated in clinical petite *vs* clinical parent (BYP41 *vs.* BYP40), such as iron-sulfur cluster assembly and transmembrane transport. The latter GO term category was also up-regulated in the clinical petite compared to a laboratory-derived one (BYP41 *vs.* D5). Finally, when comparing clinical vs non-clinical non-petite strains (BYP40 *vs.* CBS138), we observed up-regulated GO terms related to oxidative stress, fatty acid beta-oxidation and various catabolic processes. Such multifaceted differences are likely due to different genetic backgrounds of the two strains and may potentially reflect intrinsically higher capacity to respond to oxidative stress and metabolic shift in CBS138, resulting in a higher intracellular growth rate (see Figure 2A).

We then investigated the transcriptional profiles of macrophages infected with different *C*. *glabrata* strains compared to uninfected macrophages. Principal component analysis (Figure 3D) showed that at the early time point of infection, macrophages infected with CBS138 showed a different response than those infected with the three other strains, which elicited a largely similar macrophage transcriptional response. In contrast, at 24 h after infection, all strains elicited a largely uniform response, with some stratification between the response to petite and non-petite strains. Further functional GO term enrichment analysis (Figure 3E) of differentially expressed genes of infected macrophages compared to the controls showed a similar pattern to that shown in the PCA – the majority of triggered biological processes by different infecting strains were common, especially at the late stage of infection, with certain strain-specific components. For example, all infected macrophages up-regulated pathways observed in response to virus challenge, response to biotic stimulus, response to decreased oxygen levels, and response to metal ions, among others, irrespective of the infecting strain. Interestingly, all strains except strain CBS138 triggered a type I interferon response in macrophages, which was shown to be of central role in combating the major *Candida* pathogens by vaginal epithelial cells (29). Of note, this pathway was up-regulated at the 24 h time-point in macrophages infected by petite isolates. We also observed that certain pathways, such as response to interleukin-1 and several ER-related processes, were up-regulated only by clinical strains (BYP40 and BYP41).

### Petite-infected macrophages induce a pro-inflammatory transcriptional program

To dissect how fungal mitochondrial function impacts the interacting macrophages at higher resolution, we directly compared the transcriptomes of macrophages infected with non-petite and petite *C*. *glabrata* strains. Interestingly, the macrophages exhibited numerous significantly differentially expressed genes based upon whether they were challenged with petite vs. the non-petite fungal strains (Figures 3F and 3G). To further specifically identify the macrophage pathways differentially regulated in a manner dependent upon the fungal mitochondrial status, we performed gene set enrichment analysis (GSEA) (30) of the petite vs. non-petite fungal-challenged macrophage transcriptomes using the “Hallmark” Molecular Signatures Database pathways (31). GSEA revealed that both the laboratory and clinical petite strains led to the induction of pro-inflammatory pathways such as the “Interferon alpha response”, “Interferon gamma response”, “TNFA signaling via NFKB” and “Inflammatory response”. It is noteworthy that in our GO term enrichment analysis, the “Type I interferon response” at 24 h was also up-regulated exclusively in petite-infected macrophages. On the other hand, “hypoxia” and “glycolysis” pathways were consistently observed to be enriched in macrophages challenged with non-petite strains (Figure 3H).

Macrophages actively responding to diverse stimuli can have a transcriptional state that can fall across a spectrum of “classically” activated M1 to “alternatively” activated M2 phenotypes (32, 33). To assess whether the transcriptomes of petite *vs*. non-petite responding macrophages resemble an M1/M2 polarized state, we performed GSEA of our fungal-challenged transcriptomes against the human M1/M2 transcriptional modules (33). Consistent with the observed pro-inflammatory signature, the petite strain-challenged macrophages showed significant enrichment of the human M1 transcriptional module. Overall, these data suggest that the mitochondrial-deficient petite strains reprogrammed the macrophages toward a pro-inflammatory transcriptional state (Figures 3F–H).

### Intracellular petites are non-responsive to echinocandins irrespective of drug concentration

Because non-growth and slow growth in bacterial pathogens are associated with higher survival upon exposure to lethal concentrations of antibiotics (1, 5, 34), we hypothesized that *C*. *glabrata* petites could better survive lethal concentrations of cidal antifungal drugs. First, we tested whether our petites were more tolerant to general stresses, such as ER stress (tunicamycin), membrane assault (SDS), cell wall stress (Congo red), and oxidative stress (H_2_O_2_). The survival of strains BYP41 and D5 (petites) and BYP40 and CBS138 (non-petite parents) were assessed quantitatively using CFU enumeration at 3-, 6-, and 24-hours post-treatment (pt). Indeed, petites showed significantly higher survival under ER and membrane stresses, whereas they showed a similar tolerance to non-petites during cell wall and oxidative stresses (Figure 4A). A higher tolerance to ER stress has already been described for other petite *C*. *glabrata* isolates (6). Next, we assessed the survival of petites and their non-petite parent strains at 3-, 6-, 24-, and 48- hours pt when treated with 8× MIC of micafungin (0.125 µg/ml) and caspofungin (0.25 µg/ml). As hypothesized, petites exhibited a significantly slower killing rate than non-petite isolates, which mimicked microbial tolerance phenotypes characterized by slow and monophasic killing (34, 35) (Figure 4B). Since *C*. *glabrata* can survive within macrophages and intracellular petites are dormant, we explored the impact of micafungin (0.125 µg/ml) and caspofungin (0.25 µg/ml) on intracellular petite and non-petite isolates. As expected, intracellular petites were not responsive to echinocandin treatment, whereas intracellular non-petites showed a significantly higher killing rate (Figure 4C). The intracellular petites were not responsive to echinocandins irrespective of concentration and showed a much slower killing rate in RPMI compared to non-petite isolates (Figure S4A). Of note, this was somewhat concentration- and time-dependent, as planktonic petites grown in RPMI showed killing rates similar to non-petites at lower concentrations of caspofungin (0.03 and 0.06 µg/ml) after 24- hours (Figure S4B). Consistent with results obtained with other petite strains, intracellular *rdm9Δ* cells were almost 100-fold more tolerant to micafungin (0.125 µg/ml) (Fig S4C). To substantiate our findings, we established an intracellular competition assay in the presence of micafungin (0.125 µg/ml), and the proportions of GFP and RFP cells were determined by flow cytometry at 3- and 24- hours pt. In agreement with our experiments on individual strains, petites had a competitive advantage and showed a significantly higher survival than non-petites (Figures 4D and 4E; Figure S5).

**Figure 4.**
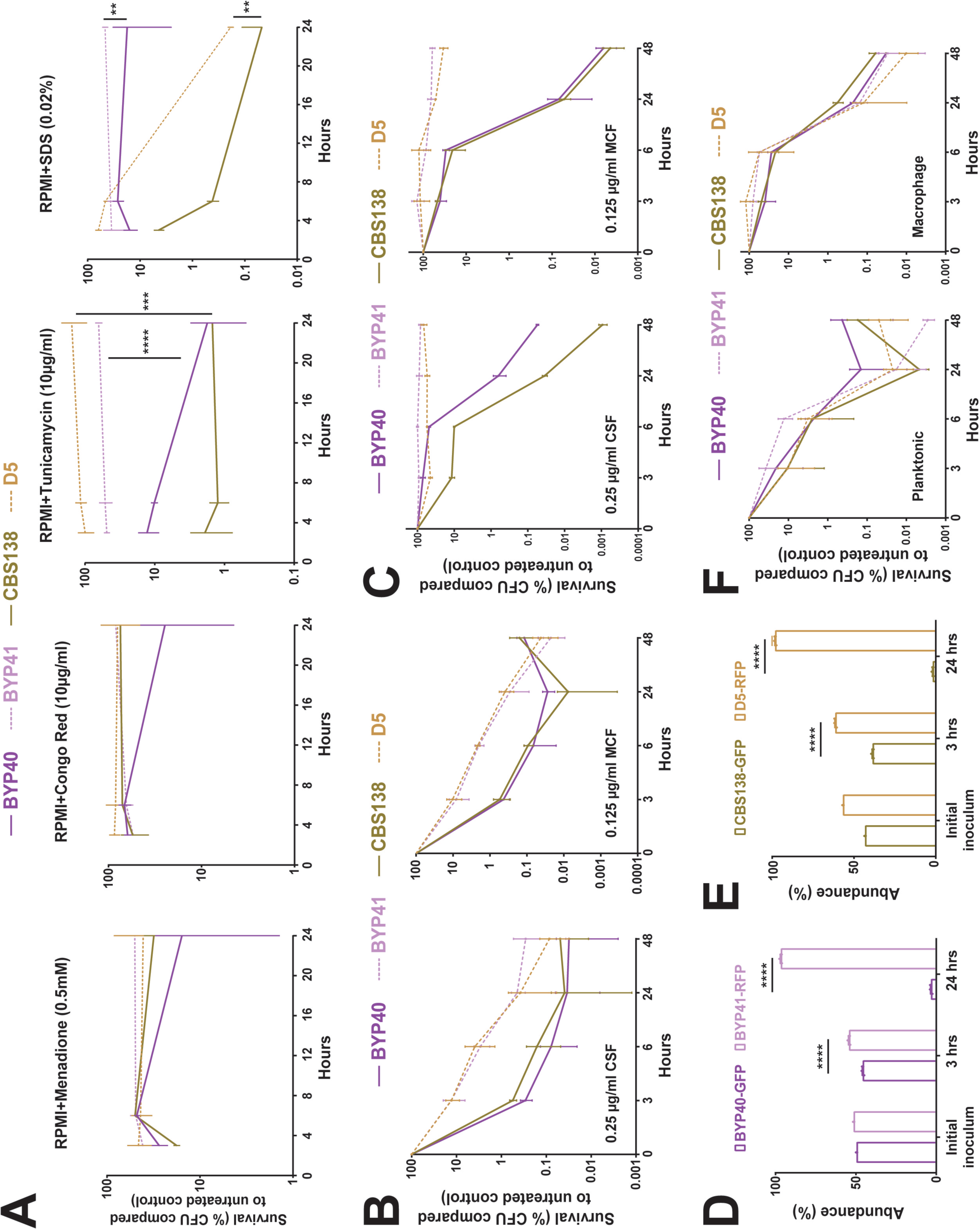
Petite phenotype is advantageous under certain stresses and echinocandin treatment. Petite and parental strains were resuspended in RPMI containing tunicamycin (endoplasmic reticulum stress; 10µg/ml), SDS (membrane stress; 0.02%), Congo Red (cell wall integrity; 10µg/ml), and menadione (0.5mM), survival was assessed at designated timepoints, and the survival data was normalized to untreated control. Petites had a higher tolerance to endoplasmic reticulum (4 biological replicates, ***<0.01 and ****<0.00001, two-tailed t-test) and membrane stresses (4 biological replicates, **=0.01, two-tailed t-test), whereas petite isolates showed similar tolerance to oxidative and cell wall stresses (all experiments were carried out in 4 biological replicates, **=0.01, ***<0.01 and ****<0.00001, two-tailed t-test) (A). The survival assessment of planktonic BYP40 and BYP41 under micafungin (0.125µg/ml) and caspofungin (0.25µg/ml) revealed that BYP41 was more tolerant and showed monophasic and slow-killing dynamic reminiscent of tolerance phenotype defined in bacteriology (8 biological replicates) (B). Intracellular BYP41 were not responsive to either micafungin or caspofungin, whereas intracellular parental strains showed 1000- to 10000-fold killing compared to petite strains (4 biological replicates) (C). Intracellular BYP40 and CBS138 were outcompeted by BYP41 and D5, respectively under micafungin treatment (3 biological replicates, ****<0.00001, two-tailed t-test) (D and E). BYP40 and BYP41 were equally killed by AMB in either planktonic (8 biological replicates) or intracellular conditions (4 biological replicates) (F).

Given that previous studies found that petites and their non-petite progenitors have similar levels of membrane ergosterol (3), the target of amphotericin B, and that the candidemia patient infected by BYP41 was successfully treated with amphotericin B (12), we reasoned that petites and non-petites should have similar killing rates by amphotericin B. In agreement with this expectation, petites and non-petites showed similar killing rates in both intracellular and planktonic conditions upon treatment with 2× MIC of amphotericin B (2 µg/ml) (Figures 4F). Although echinocandins and amphotericin B are both cidal antifungal drugs, echinocandins’ cidality requires actively growing cells producing a cell wall, whereas amphotericin B indistinguishably kills growing and non-growing cells (36). Altogether, these observations suggest that petites have a fitness benefit relative to non-petites when exposed to echinocandin drugs but are killed as effectively as non-petite cells by polyene amphotericin B.

### Petites are outcompeted by their non-petite counterparts in gut colonization and systemic infection mouse models

The non-growing phenotype of intracellular petite isolates suggested that they may be outcompeted *in vivo*. To test this hypothesis, we performed *in vivo* competition experiments between petite and non-petite strains using gut colonization and systemic infection models (37). The fecal samples collected from the gut colonization model and the kidney and spleen collected from systemically infected mouse at multiple time-points post colonization or infection, respectively, were spread on YPD plates, and the resulting colonies were visualized by a Typhoon Laser Scanner (Cytiva). Because Typhoon cannot distinguish RFP from GFP, the *in vivo* competition experiments used non-fluorescent petite and GFP-expressing non-petite isolates.

Gut colonization murine models utilized CF-1 immunocompetent mice, whose commensal gut bacteria were eradicated by piperacillin-tazobactam (PTZ) and *C*. *glabrata* colonization was induced by oral gavage as we described previously (37). Fecal samples collected at days 1, 3, 5, and 7 post colonization (pc) were plated on YPD plates containing PTZ. Mice colonized with the combination of GFP-expressing CBS138 and non-fluorescent CBS138 isolates revealed that GFP-expressing cells carry a minor fitness cost in the gut (Figure 5A). Nevertheless, the gut colonization model showed that GFP-expressing non-petite BYP40 and CBS138 outcompeted the non-fluorescent petite isolates BYP41 and D5, respectively (Figures 5B and 5C).

**Figure 5.**
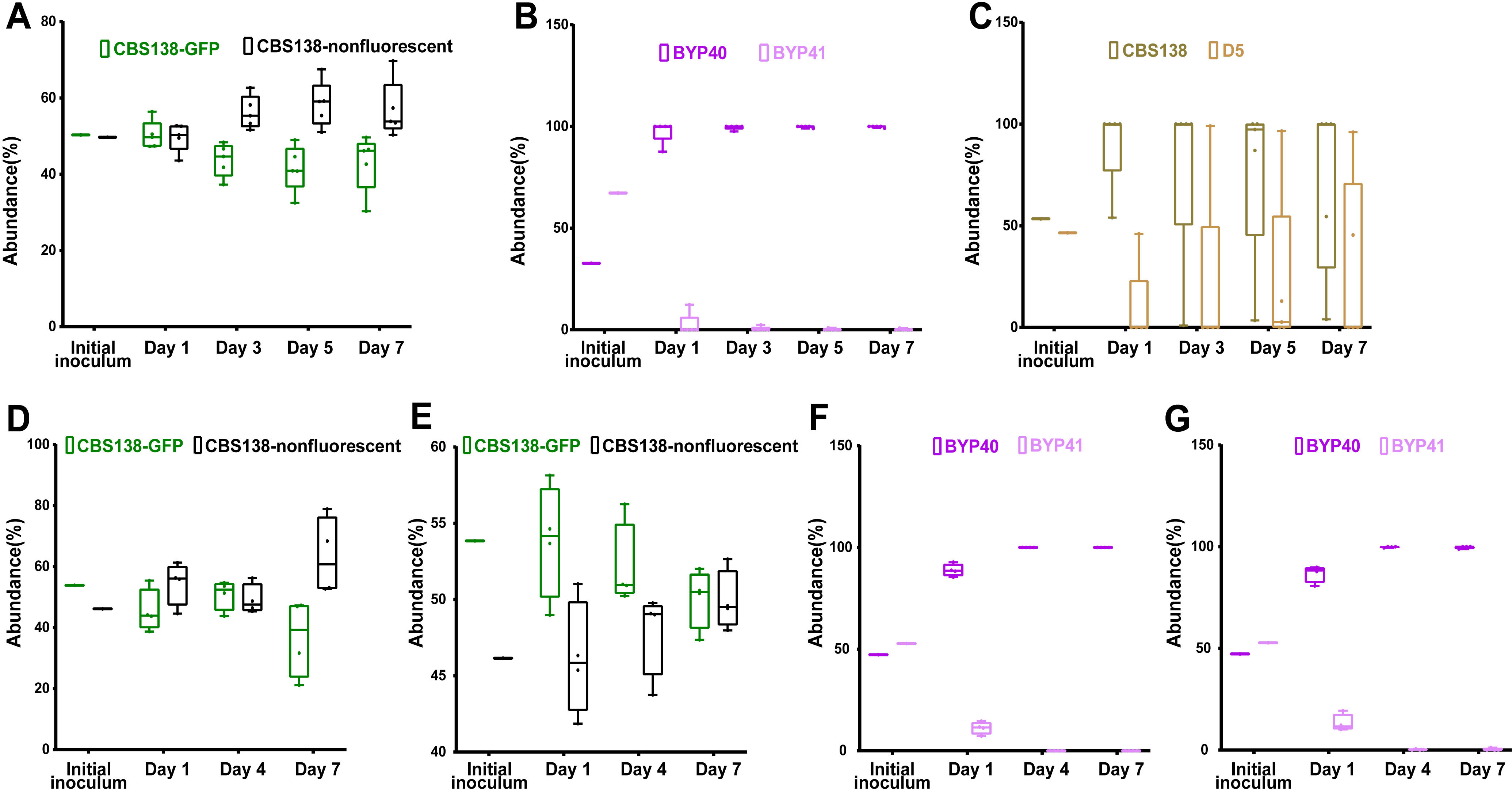
In-vivo competition of petite and their parental strains in gut colonization and systemic infection mice models. The fecal samples of CBS138-GFP and non-labelled CBS138 collected at days-1, 3-, 5-, and -7, plated on agar YPD plates, and the number of GFP- and non-fluorescent colonies were enumerated. Our gut colonization model showed a slight fitness cost of genomic GFP integration in the context of gut colonization (5 mice per each group, each dot represent one mouse) (A). Both BYP41 (B) and D5 (C) were outcompeted by their parental strains in the context of gut colonization models. Genomic GFP integration carried a significant fitness cost at day 7 in kidney (D), whereas it did not impact fitness in spleen (4 mice per each time-point, each dot represent one mouse) (E). Similar to gut colonization, BYP41 was outcompeted by BYP40 in both immunocompromised (F and G) and immunocompetent mice (H and I).

The systemic infection model utilized CD-1 female mice immunosuppressed using cyclophosphamide. The kidneys and spleens collected at 1-, 4-, and 7- days pi were homogenized and plated on YPD plates. First, we assessed the fitness cost of GFP alone in this model using non-fluorescent and GFP-expressing CBS138. Although GFP-expressing CBS138 had a fitness cost at day 7 in the kidney (Figure 5D), both GFP-expressing and non-fluorescent isolates were equally abundant in the spleen (Figure 5E). Consistent with the gut colonization results, petites were outcompeted by non-petites in both the kidney and spleen (Figures 5F and 5G). Finally, we assessed the competition of petites and non-petites in immunocompetent mice. Although petites had a higher persistence in immunocompetent mice, they were readily outcompeted by non-petite counterparts in both the kidney and spleen (Figures S6A and S6B). Altogether, these results indicated that petites are less fit relative to non-petites in the context of gut colonization and systemic infections.

### Petites show an improved survival rate *in vivo* in systemic infection mouse models during echinocandin treatment

The survival advantage of petites during echinocandin exposure in both monoculture and competition assays prompted us to ask if they can also outcompete their non-petite parental cells in systemic infection mouse models during treatment with humanized doses of caspofungin (5 mg/kg). Because petites are readily outcompeted in systemic infection mouse models, we started the caspofungin treatment either 2- hours prior (pri) to infection or 4- hours pi. Similar to our previous mouse experiments, we infected mice with an inoculum containing GFP-expressing non-petites and non-fluorescent petites, collected kidneys and spleens at days 1, 4, and 7 pi, and plated them on YPD agar plates. Although petites were again outcompeted by non-petite cells (Figures 6A–6D), their survival was significantly higher compared to the untreated condition (Figures S6A and S6B). This suggests that following caspofungin exposure, echinocandin-susceptible non-petite cells are killed more effectively than echinocandin-tolerant petite cells. These observations imply that petites have a survival advantage under echinocandin treatment *in vivo*.

**Figure 6.**
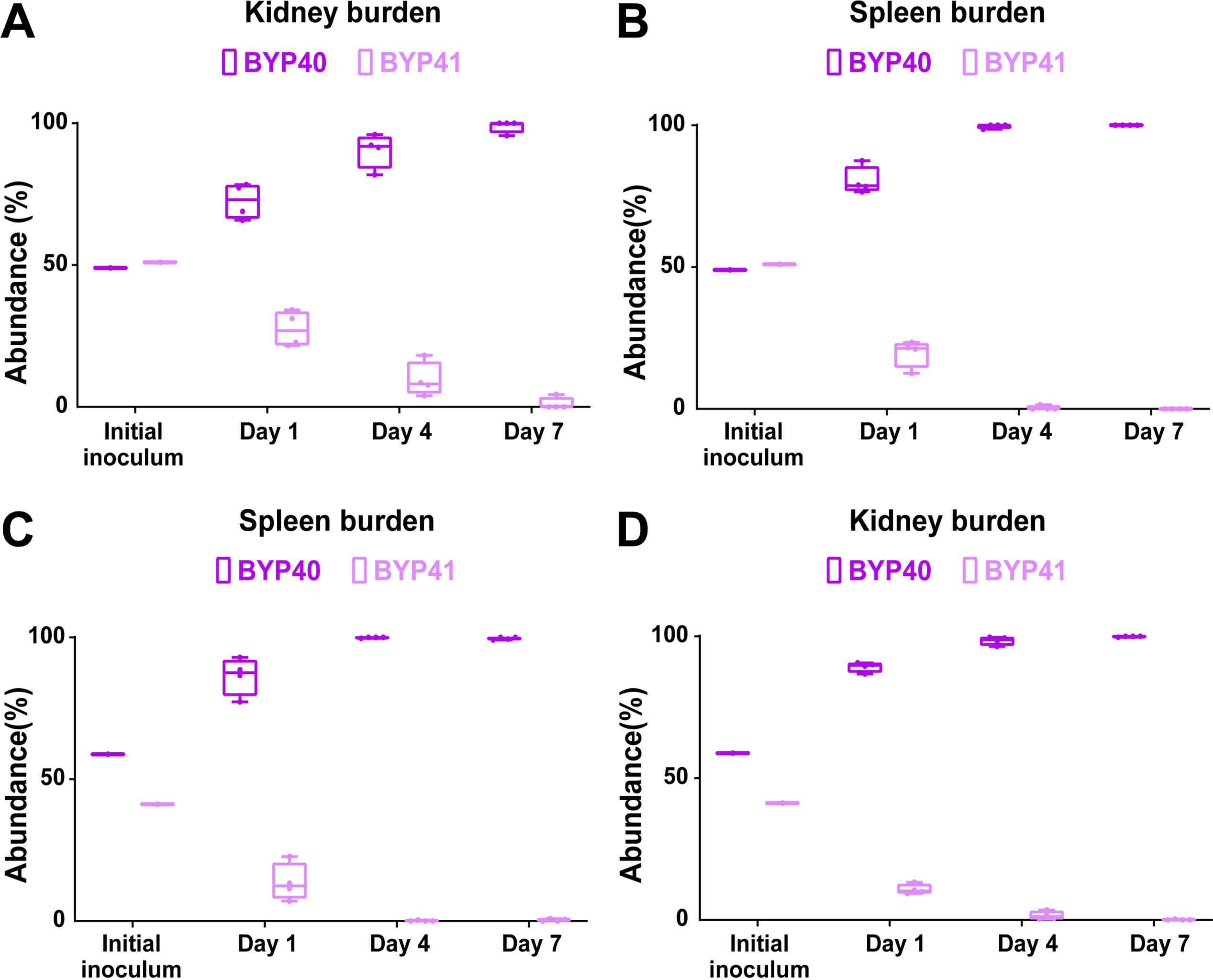
BYP41 shows an improved survival in systemic infection mice treated with humanized dose of caspfungin (5mg/kg) administered 2-hours prior (A and B) or 4-hours post-infection (C and D).

### Petite phenotype prevalence in blood isolates is rare but varies by geography

Thus far, our studies have identified the petite phenotype as relatively unfit *in vivo* in the absence of drug (echinocandins) exposure. These observations suggested that petite strains should be rare among clinical isolates of *C*. *glabrata*. Accordingly, we conducted a retrospective international study and collected >1,000 *C*. *glabrata* bloodstream isolates from Austria (*n*=600), Turkey (*n*=260), and Pakistan (*n*=220). All isolates were streaked on YPG agar plates, and those that failed to grow were considered petite. Interestingly, petite isolates were identified as pure petite colonies (*n*=1; 0.16% from Austria) and as mixtures containing both petite and non-petite colonies (*n*=8; 3.3%). It should be noted that isolates were obtained from historic glycerol stocks and authors are not sure if such isolates were obtained from a single or multiple colonies of the same sample. The pure petite blood isolate was obtained from an azole-naïve patient suffering from pancreatitis, and 62.5% (5/8) of the patients infected with the mixed isolates were not treated with azoles. This epidemiological finding suggests that although petite isolates are rarely recovered from blood samples, their prevalence varies depending on the geographical location, which may be linked to clinical practices. Moreover, these data also suggest that similar to SCV, petites can also be presented as a mixture of small (petite) and large (non-petite) colonies (4).

## Discussion

A hallmark of many yeasts is the ability to grow in the presence or absence of oxygen and form stable respiratory-deficient “petite” cells (10). The bloodstream pathogen *C*. *glabrata* can generate such petites, but their biology is only partially understood, and there has been some controversy concerning their clinical relevance. Whereas different genetic alterations can potentially cause the petite phenotype, we show that petites with different underlying mutations manifest similar responses to host and other stresses, such as the inability to proliferate inside macrophages and extreme tolerance to fungicidal echinocandin drugs. We show that in mice, petites poorly colonized the gut and poorly induced systemic infections. WGS data identified petite-specific mutations in genes mostly enriched in mitochondrial functions, which potentially underlie the phenotype, as shown here for one of them. Our dual RNAseq analysis revealed that petite-infected macrophages prominently induce pathways typically associated with type-I interferon signaling, as well as proinflammatory cytokines, and sustain these at 24 hrs compared with their non-petite counterparts. Finally, screening of a large collection of *C*. *glabrata* blood isolates from three different countries revealed that petite prevalence can vary between 0–3.3% of the total blood isolates and that similar to SCVs, petites can also be found as a mixture of small and large colonies.

Previous findings have linked the occurrence of mutations in *MIP1* to the petite phenotype in *C*. *glabrata* (7). Such mutations were not observed in our isolates, indicating that alternative mechanisms underlie the phenotype. WGS data of evolved petites identified that newly arisen mutations were enriched in mitochondria-related pathways, suggesting that mitochondrial defects drive the petite phenotype. The underlying genetic drivers are complex, and the contribution of multiple genes to the petite phenotype is yet to be fully determined. Our experiments confirmed that mutations in *RDM9* can result in a petite phenotype, underscoring the relevance of mitochondrial mRNA stability to the petite phenotype. Future studies using a wider range of clinical strains are needed to catalog a more comprehensive set of genes implicated in petite phenotype.

Similar to previous observations, we found that petite isolates had significantly higher macrophage phagocytosis rates, which may be attributed to their decreased level of β-1,3-d glucan and a compensatory increased level of mannan (7). However, contrary to previous observations (7), we found that petite isolates did not exhibit higher survival or replication inside macrophages than their non-petite counterparts, which efficiently replicated inside macrophages at 24 h. This observation was consistent with data from dual RNAseq analysis, which identified enriched over-expression of pathways involved in DNA replication and cell cycle progression in non-petite isolates. Interestingly, this *ex vivo* observation was in line with *in vivo* competition experiments, where petites were readily outcompeted by non-petites in gut colonization and systemic infection murine models. The incompetence of petite isolates in the context of *in vivo* infection models aligns with previous observations (15, 17). Although mitochondrial activities are expected to decline during the early hours of host cell infection (7, 38), our *in vivo* observations support mitochondrial function as essential to establish colonization and sustain systemic infection. Immune cells, especially macrophages, are hexose deprived, and alternative carbon sources as well as fatty acids are more abundant (38, 39). Given that oxidation of both of these types of compounds requires active mitochondria, petites are not able to assimilate such molecules to sustain intracellular growth. Moreover, RNA-seq data suggest that macrophages induce amino acid starvation (22), which would be expected to further hinder the intracellular growth of petites, given the importance of mitochondrial pathways for amino acid production (10, 18). As such, in the absence of antifungal selection pressure, petites constitute an unfit phenotype, unable to sustain host colonization and systemic infections, highlighting the critical role of mitochondrial functions to successfully colonize mucosal surfaces and cause systemic infections. In contrast, Ferrari et al. observed that BYP41 was able to outcompete BYP40 in the context of systemic infection and vaginal colonization mouse models (14). Although further studies are required to elucidate the reason underlying this controversy, we believe that it may have stemmed from technical variations in determining the ratio of petite/non-petite from clinical samples. Given that petite isolates are fluconazole resistant, but not the non-petite parental strains, Ferrari et al. plated homogenates on YPD plates containing 30 µg/ml of fluconazole to discern the proportion of petite over non-petite (14). Nonetheless, in our study, both petite and non-petite isolates were able to grow on YPD plates containing 30 µg/ml of fluconazole after 48 hrs of incubation. Therefore, we generated fluorescently-labelled strains to overcome this obstacle and to accurately measure the ratio of petite/non-petite colonies for our *in vivo* mouse models. Our results are consistent with a previous study, where authors noted that mice infected with petite had a significantly lower fungal burden compared to non-petite across all the organs tested (15). Consistently, petite *C*. *glabrata* colonies isolated from blood samples from a recent study were also efficiently cleared in the spleen, liver, and kidney of a systemic infection mouse model (40).

Bacteria can adopt various phenotypes, such as SCV (1, 4, 8) and antibiotic tolerant and persister cells (34), which are not effectively killed by cidal antibiotics or by other host-related stresses. Such phenotypes are marked by lower metabolic activities and by arrested or slow growth (1, 4, 34, 41). When exposed to antibiotics, tolerant bacteria display slowed and monophasic killing curves owing to specific genetic mutations that exist throughout the clonal population, whereas persisters are characterized by biphasic killing and do not carry any genetic changes (34). Interestingly, upon exposure to caspofungin and micafungin, petite isolates displayed higher tolerance and a monophasic killing pattern. In addition, similar to antibiotic tolerant cells, petite isolates harbored mutations in multiple genes potentially underpinning echinocandin tolerance. The parental strains, however, showed the hallmark of echinocandin persisters, as described previously (36). Of note, both phenomena are distinct from the concept of azole tolerance, which refers to slow growth in the presence of static antifungals (42), whereas echinocandin tolerance and persistence refer to survival during exposure to supra-MIC concentrations of cidal antifungal drugs. Interestingly, due to their lack of intracellular growth, the tolerance level of petites was significantly increased after phagocytosis, and accordingly, intracellular petites outcompeted the parental strains after exposure to cidal concentrations of echinocandins. *In vivo*, petite isolates were outcompeted by a mitochondrial proficient parental strain. However, following drug exposure, petites had a higher survival in mice treated with humanized dosage of caspofungin when compared to untreated control mice. How petites can cause systemic and superficial infections and outcompete their mitochondrial proficient kins in humans is currently unknown. Petites may need certain biological niches in the host, which requires time to be established, and these factors could have been missed in a murine model of acute systemic infection. In fact, petites are not considered an end-stage phenotype, as they can convert to mitochondrial proficient cells under certain *in vitro* (10) and in-host (38) conditions. The potential physiological and clinical relevance of this observation needs to be investigated further.

In agreement with the general observations that both petite and non-petite isolates appear to have similar levels of ergosterol on the cell membrane (3) and the observation that the patient infected with BYP41 was successfully treated with amphotericin B (12), our *in vitro* and *ex vivo* analyses found similar killing efficiency for petite and non-petite isolates treated with amphotericin B. This similar killing rate is at least partly due to the observed lack of growth dependence of amphotericin B compared to echinocandins (36). Although additional *in vitro* and clinical trials are needed, our experiments suggest that amphotericin B treatment may be especially suitable for patients chronically infected with *C*. *glabrata*.

Because the implications of the petite phenotype for interactions with host cells have remained elusive, we compared the transcriptomic responses mounted by macrophages infected by petite and non-petite *C*. *glabrata*. We noted that petite-infected macrophages mounted a more pronounced type-I interferon (TII) response and marked overexpression of genes associated with proinflammatory cytokine signaling at 24 hours pi. Although classically associated with viral infection, this response is also induced by bacterial, parasitic (43) and fungal infections (29, 44–48). Similar to bacterial infections, the TII response appears to play a somewhat controversial role in fungal infections, which varies depending on the host cell type and the fungal species (29, 44–48). Indeed, it has been noted that the TII response is beneficial for the survival of *C*. *glabrata* cells in macrophages (45, 48), and accordingly, it is plausible to assume that the TII response mounted by macrophages provides a permissive environment for the long-term survival of petites in the host. Intriguingly, host cell mitochondrial DNA release at early hours pi promotes a TII response in vaginal epithelial cell lines infected with *Candida* species (29). Although our experiments were performed in a different cell type, it could be speculated that the late TII response in petite-infected macrophages results from the reduced damage inflicted upon macrophages by dormant petite cells, as supported by our cellular damage assays. Future studies are warranted to uncover the stimuli behind such a response and whether inhibition of the TII response in macrophages could decrease the burden of petites. The overexpression of genes associated with cytokine production was another hallmark of petite-infected macrophages. This transcriptomic rewiring is potentially driven by cell wall carbohydrate differences (7) leading to differential C-type lectin receptor signaling. Since hyper-inflammation in a given niche is associated with pathological development (49), future studies are warranted to show whether infection with petite *C*. *glabrata* is associated with any currently unknown pathological manifestations.

Strikingly, petites are more abundant in urine samples (7, 12, 17), and a recent study identified 10.2% (15/146) petite isolates in a collection of *C*. *glabrata* clinical isolates from diverse clinical samples (7). The prevalence of petite isolates also appeared to be 10% (1/10) in a recent candidemia study exploring the genotypic diversity of *C*. *glabrata* colonies growing from the same blood sample (40). Although our retrospective, multicenter, international study assessing >1,000 *C*. *glabrata* blood isolates identified only 9 petite isolates (0.9%), the petite prevalence varied depending on the country and ranged from 0 to 3.3% of the total isolates. Interestingly, all of the petite isolates identified in Turkey were a mixture of small and large colonies. Although this may reflect the differences in clinical practice, it is also possible that some genotypes may have a higher propensity to develop petite colonies. It is noteworthy that the clinical isolation procedure may inadvertently select against slow-growing colonies in mixed populations, effectively rendering petite cells undetected and thus underestimating petites as a clinical entity. As such, our study suggests that the identification of petite *C*. *glabrata* isolates and appreciation of their clinical relevance requires a comprehensive and unbiased characterization of both small and large colonies. In keeping with this hypothesis, a recent study showed that petite colony recovery from blood cultures required ≥84 hours of incubation to become evident on agar plates, and as such, they went undetected by the microbiology laboratory. Interestingly, given that the index colony tested was azole susceptible, the patient was treated with fluconazole, and this underestimation of petite resulted in fluconazole therapeutic failure and the emergence of pure petite isolates from later blood samples (40). Therefore, the microbiology laboratory practices and the slow growing nature of petites from blood (and maybe other sterile samples) may have resulted in underestimation of petite phenotype. This phenotypic plasticity and the switching from large to small colonies and *vice versa C*. *glabrata* to effectively linger in the host and adopt various metabolic states depending on the host conditions. Accordingly, depending on the proportion of small and large colonies during the course of infection, one may be able to predict the therapeutic efficacy of antifungal drugs.

Petite isolates are known to emerge following fluconazole treatment (3, 12), yet the majority of petite-infected patients are azole-naïve (7, 17). This is consistent with the previous observation that petite *C*. *glabrata* isolates also emerge following macrophage internalization (7). Therefore, prospective, observational studies employing a wide range of clinical samples may more accurately reflect the prevalence of petites in the clinic and potentially discover inducers that could drive petite emergence.

## Methods

### *C*. *glabrata* strains and growth conditions

Our clinical collection included 37 isolates, two of which were petite (BYP41 and DPL248) and the rest were non-petite. Moreover, four isogenic petite isolates, C5, D5, F2, and G5, were derived from CBS138 (Supplementary Table 1). Except for BYP40, BYP41, DPL248, and CBS138 and petite mutants derived from it, the rest of the isolates were only used for MIP1 sequencing. Details regarding the generation of petite isolates are described in the results. All isolates were grown on YPD agar and broth overnight. YPD contained 10 g/L of yeast extract, 20 g/L of peptone, 20 g/L of dextrose, whereas in YPG, dextrose was replaced with 20 g/L of glycerol.

### Macrophage infection

We used human THP1 macrophages derived from the human acute monocyte leukemia cell line (THP1; ATCC; Manassas, VA, USA). THP1 macrophages were grown in RPMI 1640 1640 (Gibco, Fisher Scientific, USA) containing 1% pen-strep (Gibco) and 10% heat-inactivated HFBS (Gibco). Two days prior to infection, one million THP1 monocytes were treated with 100 nM phorbol 12-myristate 13-acetate (PMA, Sigma) in 24-well plates and incubated in a 5% CO_2_ incubator at 37°C. On the day of infection, macrophages were washed with PBS and treated with fresh RPMI 1640. Overnight-grown *C*. *glabrata* cells were washed three times with PBS, and depending on the experiment, the macrophages were infected with a multiplicity of infection (MOI) of 10 *C*. *glabrata*/1 macrophage, 1/1, or 10/1. Given that *C*. *glabrata* can grow up to 5- to 7-fold inside macrophages after 24 hours and to accurately measure the intracellular growth rate, we used an MOI of 1/10, whereas experiments studying the cytotoxicity and impact of intracellular killing by antifungals used MOI of 1/1 and 10/1, respectively. Of note, untreated control macrophages were again infected with MOI of 1/10 given that the impact of antifungal drugs was assessed for up to 48 hours, and a dilution factor of 100 was considered in intracellular survival determination. After a 3-hour incubation, the extracellular, non-adherent *C*. *glabrata* strains were extensively washed with PBS and treated again with fresh RPMI. The first wash was plated on a YPD plate to measure the phagocytosis rate. Macrophages were lysed by the addition of ice-cold water and extensive pipetting to effectively lyse the macrophages and release the intracellular *C*. *glabrata* (ICCG) cells, and the lysates were transferred into YPD agar plates.

### Petite determination

Potential petite isolates unable to grow on YPG agar plates were subjected to the following experiments to ensure that such isolates were *bona fide* petite mutants. ATP was determined using a luciferase kit (Thermo Fisher). Briefly, exponentially growing cells in RPMI were washed twice with PBS, and the pellets were incubated in Y1 buffer containing 100 units of lyticase (Sigma) at 37°C for 30 minutes. Subsequently, the pellets were resuspended in ATP extraction buffer (Thermo Fisher), incubated at 80°^°^C for 10 minutes, and subjected to bead-beating for two minutes. Finally 20 µl of these samples were added to master mixes containing luciferase, and the luminescence was measured using a plate reader. Mitochondrial membrane activity was measured using rhodamine 1,2,3 (Sigma) from exponentially growing *C*. *glabrata* cells using flow-cytometry (BD Bioscience). The basal expression levels of *CDR1*, *CDR2*, *PDR1*, and *SNQ2* were determined from exponentially growing *C*. *glabrata* using the primers listed in Supplementary Table 4, second sheet and the expression values were normalized using RDN5.8 primers described elsewhere (6). RNA was extracted using a previously described method (29), followed by DNase treatment (QIAGEN) and repurification of RNA samples using the QIAGEN RNeasy Kit. Of note, dual RNA-seq analysis as well as the RNA samples used to determine the expression levels of *FKS1* and *FKS2* used the same RNA extraction methodology. Fluconazole resistance was determined when *C*. *glabrata* cells showed an MIC≥64 µg/ml following the Clinical Laboratory Standard Institute procedure, and the MICs were determined visually and using a plate reader.

### Metabolite dependency determination

To determine the impact of metabolites on the growth rate of petite and non-petite isolates, we grew *C*. *glabrata* cells overnight in YPD broth and then washed twice with PBS. The OD was adjusted to 0.1, 200 µl of cell suspension was placed in a 96-well plate sealed with a breathable film cover (Sigma), and the kinetic growth rate was monitored using a Tecan plate reader for 16- hours. The final growth of YNB individually supplemented with arginine (20 mg/L), leucin (60 mg/L), glutamine (2 mM), glutamate (5 mM), aspartate (20 mg/L), menadione (5 µg/ml), thymidine (100 µg/ml), adenine (20 mg/L), and hemin (1 µg/ml) was subtracted from that of YNB alone, and the average values were used to draw a heatmap using an on-line free tool.

### Gut colonization mouse model

We used a previously established GI-tract mouse model (37), which uses 6-week-old female CF-1 mice (Charles River Laboratory). To effectively establish *C*. *glabrata* colonization and eradicate commensal gut bacteria, mouse were subcutaneously treated with piperacillin-tazobactam starting 2 days prior to infection and continued daily until the end of the time-course. Colonization was induced by oral gavage using 1.5 × 10^8^ cells in 100 µl of sterile PBS. Fecal samples were collected on days 1-, 3-, 5-, and 7-post colonization and 100 µl of fecal samples were plated on YPD plates containing PTZ. Plates incubated at 37°C for up to 2 days, and plates were visualized using Typhoon imaging.

### Systemic infection mouse model

Systemic infection used six-week-old CD-1 mice (Charles River Laboratory), which were immunosuppressed with cyclophosphamide starting from 3 days prior to infection (150 mg/kg) and continued every 3 days once with 100 mg/kg (37). Infection was established via the rhino-orbital route using 50 µl of cell suspension containing 5 × 10^7^ cells. Kidney and spleen samples collected on days 1-, 4-, and 5-pi were homogenized, 100 µl of which was streaked on YPD agar, incubated for up to two days in a 37°C incubator, and visualized using Typhoon.

### Knockout generation

Deletant mutant *C*. *glabrata* colonies were generated using previously described methods (6). Briefly, the nourseothricin cassette containing flanking regions homologous to the outside ORF of the desired genes was amplified using the primers listed in Supplementary Table 4, which were used for transformation. Competent *C*. *glabrata* cells were created using a Frozen-EZ Yeast Transformation Kit (Zymo Research) and followed an electroporation-based methodology described previously. Transformed yeast cells were transferred onto NAT-containing YPD, and colonies were subjected to diagnostic primers listed in Supplementary Table 4.

### *MIP1* and *PDR1* sequencing

*PDR1* sequencing followed previously described primers and PCR conditions (50), and the primers used to amplify and sequence *MIP1* are listed in Supplementary Table 4.

### Macrophage damage assay

To measure the extent of damage incurred by *C*. *glabrata* isolates to macrophages, we measured the level of lactate dehydrogenase using a commercial kit (Sigma) (29). Briefly, macrophages infected with an MOI of 5/1 were extensively washed with PBS 3 hr pi, followed by the addition of fresh RPMI and incubation in a CO_2_ incubator at 37°C for another 21 hr. After 24 hours, supernatant samples were collected, and LDH was determined as described previously (29). The OD value of each replicate was subtracted from that of the background control (uninfected macrophages), and the corrected value was divided by that of high control (uninfected macrophages treated with 0.25% Triton X-100 for 3 minutes). The corrected normalized values are presented as percentages.

### RNA extraction

Macrophages infected with an MOI of 5/1 were extensively washed 3 hr pi, and fresh RPMI was added to the wells to be further incubated at 37°C. After extensive PBS wash at each step, macrophages were subjected to a manual RNA extraction protocol described elsewhere (29). The RNA samples were treated with RNase free-DNase and further purified using an RNeasy kit (QIAGEN) per the manufacturer’s instructions. The integrity and quantity of RNA samples were confirmed by running RNA samples in 1.5% agarose gel and NanoDrop (Thermo Fisher), respectively.

### RNA-seq

RNA samples were quantified using Qubit 2.0 Fluorometer (Life Technologies, Carlsbad, CA, USA), and RNA integrity was checked using Agilent TapeStation 4200 (Agilent Technologies, Palo Alto, CA, USA). The RNA sequencing libraries were prepared using the NEBNext Ultra II RNA Library Prep Kit for Illumina according to the manufacturer’s instructions (New England Biolabs, Ipswich, MA, USA). Briefly, mRNAs were initially enriched with oligo(T) beads. Enriched mRNAs were fragmented for 15 minutes at 94°C. First strand and second strand cDNA were subsequently synthesized. cDNA fragments were end repaired and adenylated at the 3’ ends, and universal adapters were ligated to cDNA fragments, followed by index addition and library enrichment by PCR with limited cycles. The sequencing libraries were validated on the Agilent TapeStation (Agilent Technologies, Palo Alto, CA, USA) and quantified by using Qubit 2.0 Fluorometer (Thermo Fisher Scientific, Waltham, MA, USA) as well as by quantitative PCR (KAPA Biosystems, Wilmington, MA, USA). The sequencing libraries were clustered on four flow cell lanes. After clustering, the flow cell was loaded on the Illumina HiSeq instrument (4000 or equivalent) according to manufacturer’s instructions. The samples were sequenced using a 2 × 150bp paired end (PE) configuration. Image analysis and base calling were conducted by the Control software. Raw sequence data (.bcl files) generated from the sequencer were converted into fastq files and de-multiplexed using Illumina’s bcl2fastq 2.17 software. One mismatch was allowed for index sequence identification.

### Genome sequencing

DNA was fragmented to sizes between 1 and 20 kb using a transposase that binds biotinylated adapters at the breaking point. Strand displacement was performed to “repair” the nicks left by the transposase. Fragment sizes of 3 to 6 kb were then selected on a 0.8% agarose gel and circularized. Non-circularized DNA was removed by digestion. The circular DNA was then mechanically sheared into fragments of 100 bp to 1 kb approx. The fragments containing the biotinylated ends were pulled down using magnetic streptavidin beads and submitted to a standard library preparation. A final size selection on a 2% agarose gel was performed, and fragments of 400 to 700 bp were selected for the final library. Final libraries were analyzed using an Agilent High Sensitivity chip to estimate the quantity and check size distribution and were then quantified by qPCR using the KAPA Library Quantification Kit (ref. KK4835, KapaBiosystems) prior to amplification with Illumina’s cBot. Libraries were sequenced 2 × 150 bp on Illumina’s HiSeq 2500.

### Genome analyses

All sequencing data were processed to call single nucleotide polymorphisms (SNPs) using Freebayes (51), HaplotypeCaller (52), and Bcftools (53) as implemented in PerSVade v 1.0 (54) and using the genome sequence of CBS138 *A22-s07-m01-r86* as a reference (55). Coverage was calculated for all strains analyzed at a chromosomal level, including mitochondria, using mosdepth (56). SNPs for which a mean mapping quality was below 30, a QUAL value was below 20 or read depth was below 30 were filtered out. High confidence variant calls (those supported by two or more callers) were considered. To identify recent variants appearing specifically in the strains with phenotype phenotype (petite-specific variants), we considered high confidence variants that were differentially called between related strains differing in this phenotype. We discarded such differences when the same variant was called with an high confidence in one strain and with low confidence in another strain with alternative phenotype. As DPL248 lacked a closely related parental strain and had many polymorphisms with respect to the CBS138 reference, we used a PubMLST search by locus to identify the closest sequence type we could use based on its allelic profile (57). The two closest matches were CST35 and EB0911Sto (20). We downloaded their raw reads (accession PRJNA361477) and ran the same variant calling pipeline. Then, we used their variants for low confidence filtering in DPL248. Additionally, we manually filtered out likely artifactual mutations through visual inspection using the Integrative Genomics Viewer (IGV) (58). As BYP40 and BYP41 were related but only the latter had the petite phenotype, only SNPs unique to each of them were considered. We also filtered out SNPs that overlapped between two or more of the three petite strains derived from CBS138 (C5, D5, F2, and G5), as these shared variants were likely present in the parental strain used in the experiments and are thus unrelated to the petite phenotype. Finally, we selected genes with petite-specific variants in two or more of the three studied clonal sets. The final list of selected SNPs was processed through Variant Effect Predictor (59) and manually curated by inspecting read alignments in the region.

### RNAseq Analysis

FastQC v. 0.11.8 (https://www.bioinformatics.babraham.ac.uk/projects/fastqc/) and MultiQC v. 1.12 (60) were used to perform quality control of raw sequencing data. Read trimming was performed using Trimmomatic v. 0.36 (61) with the following parameters: TruSeq adapters: 2:30:10 LEADING:3 TRAILING:3 SLIDINGWINDOW:4:3 MINLEN:50.

For read mapping and quantification, we used the splice junction-sensitive read mapper STAR v. 2.7.10a (62) with default parameters. For samples comprising exclusively either fungal or human RNA, reads were mapped to the corresponding reference genomes. For samples containing RNA from both host and pathogen, reads were mapped to the concatenated human and yeast reference genomes. For human data, we used the novel T2T CHM13v2.0 Telomere-to-Telomere genome assembly (63) genome annotations from the NCBI (last accessed on 12 May 2022). This assembly lacked mitochondrial DNA, and therefore, we added the human mitochondrial genome of the GRCh38 human genome assembly obtained from the Ensembl database (last accessed on 12 May 2022, (63)). Reference genomes and genome annotations for *C*. *glabrata* CBS138 were obtained from the Candida Genome Database (CGD, last accessed on 12 May 2022, (55)). Potential read-crossmapping rates (i.e., reads that mapped equally well to both human and fungal genomes) were assessed with crossmapper v. 1.1.1 (55). Further downstream analyses were carried out with R v. 3.6.1. Differential gene expression analysis was performed using DESeq2 v. 1.26.0 (64). Genes with |log-2 fold change (L2FC) | > 1 and adjusted p value (padj) < 0.01 were considered differentially expressed. Gene Ontology (GO) term enrichment analysis of differentially expressed genes and enrichment visualization were performed by ClusterProfiler v. 3.14.3 (65). For GO term enrichment analysis, we used the “Biological Process” category. GO term association tables for *C*. *glabrata* were obtained from CGD (last accessed on 12 May 2022), whereas for human data, we used genome-wide annotation for the Human (i.e., org.Hs.eg.db) database v. 3.10.0 to perform GO enrichment tests.

### Gene set enrichment analysis

To define differentially enriched pathways in petite *vs*. non-petite challenged THP-1 macrophages, we utilized DESeq2 normalized counts and performed gene set enrichment analysis (30) with GSEA version 4.2.3. We tested enrichment of the “Hallmark” Molecular Signatures Database pathways (31) where weighted enrichment statistic was used with “Signal2Noise” as a metric for ranking genes. The gene sets with a false discovery rate (FDR) of less than 0.05 was determined. To assess the enrichment of M1/M2 transcriptional modules, GSEA was performed using the gene sets (transcriptional modules) described by Xue et al. (33).

## Data availability

Raw sequencing data were deposited in the Sequence Read Archive (SRA) database under project accession numbers PRJNA901527 (for RNA-seq) and PRJNA901678 (for WGS).

## Acknowledgement

This work was supported by NIH 5R01AI109025 to D.S.P.

## Figure Legends (Supplemental)

**Figure S1.** Describes the processes involved in the development of laboratory and clinically derived petite *C*. *glabrata* isolates. The petite mutant isolates recovered from CBS138 were obtained under the selection pressure of fluconazole (A). BYP41 was derived from an immunocompetent patient after fluconazole treatment and was genetically close to BYP40 (B). Supplementation with certain metabolites fosters the growth rate of petite isolates. Petites grow poorly on yeast-nitrogen-based (YNB) media, reflecting their defective mitochondria and their inability to assimilate non-fermentable carbon sources. To determine the metabolic deficiencies of *C*. *glabrata* petite strains, we measured their growth rates in yeast nitrogen base (YNB) medium and YNB individually supplemented with arginine (20 mg/L), leucine (60 mg/L), glutamine (2 mM), glutamate (5 mM), histidine (20 mg/L), lysine (60 mg/L), aspartate (20 mg/L), menadione (5 µg/ml), thymidine (100 µg/ml), adenine (20 mg/L), and hemin (1 µg/ml). Consistent with this, BYP41 and C5, D5, and F2 were more similar in their metabolite dependency profiles, whereas G5 and DPL248 clustered with non-petite isolates. The addition of leucine, arginine, and glutamine significantly improved the growth rate of petite isolates (C). As expected, petites grew slowly in unsupplemented YNB, with G5 and DPL248 growing better than BYP41, C5, D5, and F2 but still more slowly than non-petite strains (D).

**Figure S2.** Mitochondrial DNA (mtDNA) coverage of petite and non-petite isolates using whole-genome sequencing identified that petite isolates have a lower mtDNA than non-petites, with BYP41 having the lowest mtDNA content (A). Similar to petite isolates, the *Rdm9Δ* isolate was resistant to fluconazole (B). Similar to petite isolates, *Rdm9Δ* overexpresses efflux pumps and *PDR1* compared to the parental strain CBS138 (C).

**Figure S3.** Similar to petite isolates, *Rdm9Δ* has a significantly lower level of ATP than the parental strain CBS138 (A). Similar to petite isolates, *Rdm9Δ* has a lower mitochondrial membrane potential than the parental strain CBS138 (B). Similar to petite isolates, *Rdm9Δ* shows poor growth on YNB (C), and supplementation with specific metabolites ameliorates its growth (D). Similar to petite isolates, the *Rdm9Δ* isolate does not grow inside macrophages.

**Figure S4.** Non-responsiveness of intracellular petites to echinocandins. Unlike non-petite parental isolates, petites are not efficiently killed by echinocandins upon engulfment by macrophages (A). Petite isolates have a higher echinocandin tolerance to various echinocandin concentrations once incubated in RPMI (B).

**Figure S5.** Flow cytometry gating and strategy used to differentiate GFP and RFP under micafungin treatment. Similar to petite isolates, intracellular *rdm9Δ* is extremely tolerant to micafungin.

**Figure S6.** Mice infected with the petite isolate BYP41 had a lower burden, especially in the kidney at early timepoints, compared to mice treated with a humanized dosage of caspofungin (A). The treated arm included mice that received caspofungin 2 hrs prior to infection or 4 hrs post-infection (B).

